# Phylogenomic placement and morphological description of a novel phagotrophic euglenid from Hawaii: *Hokulea waialensis* n. gen. et sp

**DOI:** 10.1101/2025.10.01.676111

**Authors:** Alfredo Rodriguez Ruiz, Maia V. Palka, Gordon Lax, Dagmar Jirsová, Gianluca Fuggiti, Yu-Ping Poh, Brian S. Leander, Jeremy G. Wideman

## Abstract

Euglenids are a diverse group of flagellated protists that include phagotrophic, osmotrophic, and phototrophic lineages. Understanding the phylogenetic relationships of phagotrophic euglenids is crucial in understanding euglenid evolution as a whole. Yet many relationships within euglenids remain unclear, and further resolution requires extensive sampling, particularly from the deep-branching, paraphyletic group known as ‘ploeotids’. Improved resolution of evolutionary relationships among ploeotid taxa is necessary to elucidate the origin and diversification of complex ultrastructural traits (e.g., pellicle, feeding apparatus). Here, we isolated, cultivated and characterized a novel ‘ploeotid’ species named *Hokulea waialensis* n. gen. et sp. using light and scanning electron microscopy, single-cell sequencing, and phylogenomic analyses. This new species is relatively small (10-12 µm long) compared to related euglenids, and shares several morphological traits with related species of Alistosa. Both single and multigene phylogenetic reconstructions from single amplified genome data show that *Hokulea waialensis* n. gen. et sp. is closely related to several environmental small subunit ribosomal DNA (SSU rDNA) sequences, and more broadly to *Lentomona*s and *Decastava*.

## Introduction

Resolving the deep phylogenetic relationships within the Euglenozoa requires adequate representation of all its major groups, especially the undersampled phagotrophic euglenids. Euglenids are a large, diverse group of flagellates that display various forms of nutrition and novel morphological characteristics, such as a complex cytoskeleton with proteinaceous pellicle strips, and a robust feeding apparatus (Leander et al. 2007, 2017; Kostygov et al. 2021). Euglenids can be divided into three major groups based on morphology (e.g. pellicle anatomy, number of visible flagella): the Spirocuta, Petalomonadida, and the paraphyletic ‘ploeotids’. Certain spirocutes are the best-studied euglenids as they contain the well-known phototrophic euglenids (e.g. *Euglena*), but also include osmotrophic and many phagotrophic species (Lax et al., 2023). Petalomonads are rigid inflexible cells with one or two flagella and form a clade that branches at the base of euglenids, whereas ‘ploeotids’ are biflagellated, rigid, and use their anterior flagellum for feeding and glide on their posterior flagellum (Leander et al., 2007, 2017). ‘Ploeotids’, known to feed on eukaryotes and/or bacteria (Lax et al., 2019; Palka et al., 2025), are paraphyletic being composed of at least two clades (Lax et al., 2020, Lax et al., 2023). Better taxon sampling has resulted in more robust single- and multi-gene phylogenetic analyses and has increased our understanding of deeper branching phagotrophic lineages (Cavalier-Smith 2016; Lax et al. 2021, 2023; Schoenle et al. 2019). However, many gaps remain in our understanding of phagotrophic euglenids.

Euglenids have several distinctive traits, perhaps the best example is the pellicle, which is composed of proteinaceous strips that run longitudinally or helically under the cell membrane (Leander and Farmer 2000; Leander and Farmer 2001; Leander 2004; Leander et al. 2007, 2017). The number of pellicle strips in euglenids varies drastically between different species, ranging from < 10 in petalomonads to 120 in some spirocutes (e.g., *Euglena obtusa*) (Leander 2004; Leander et al. 2007; Esson and Leander 2008). In euglenids with > 20 pellicle strips, the strips are supported by an underlying system of microtubules that facilitates euglenoid movement (Leander et al., 2001.; Leander and Farmer 2001; Leander 2004; Leander et al., 2007, 2017; Yubuki and Leander 2012). Euglenids with < 12 pellicle strips, like petalomonads and ‘ploeotids’, have rigid cell shapes (Leander et al., 2017).

The backbone phylogeny of phagotrophic euglenids is only beginning to be resolved. Although petalomonads form a well-supported clade in phylogenies inferred from both small subunit ribosomal DNA (SSU rDNA) sequences and multigene datasets, the branching order of petalomonads and the various groups of ‘ploeotids’ (e.g., Alistosa and Karavia) remains unresolved, even in multigene phylogenies (Lax et al 2023). Thus, more sampling of these groups is required. Analyses focused on SSU rDNA sequences have expanded the taxon representation for phagotrophs in single-gene trees have helped shed light on complex relationships between these species. However, the limitations of single-gene trees, along with the fact that euglenid SSU rDNA varies so much in size and is extremely divergent, have complicated these analyses (Lax et al., 2023, Lax and Simpson 2020, Lax and Simpson 2013). Multigene analyses have strengthened our understanding of the phylogenetic relationships between various taxa, though many branches remain unresolved (Lax et al., 2021,Lax et al., 2023).

Here, we describe *Hokulea waialensis* n. gen. et sp., a new ‘ploeotid’ genus from Hawaii, using both light and scanning electron microscopy. We also use single-cell genomics, SSU rDNA phylogenetics, and phylogenomic analyses to demonstrate that *H. waialensis* n. gen. et sp. branches within the ploeotid group Alistosa close to genera like *Decastava* and *Lentomonas*. These data improve the resolution of deep branching patterns within phagotrophic euglenids.

## Materials and Methods

### Sampling, isolation, and cultivation

*Hokulea waialensis* was collected by scraping rocks near Wai’ale Falls on Wailuku River (19.715° N, −155.138° W) on the island of Hawaii near Hilo in 2021. Samples were left in the dark for ~ 2 weeks to eliminate photosynthetic organisms. Euglenids were initially identified under bright field microscopy at 400x magnification. A mono eukaryotic culture was established via single-cell isolation into a ventilated cell culture flask containing freshwater DY-V media (Andersen 2009), and kept at 23°C. Cultures were sub-cultivated weekly in a 1:1 ratio fresh DY-V medium to original culture.

### Differential interference contrast microscopy (DIC)

Specimens were further examined under and recorded using a Nikon (Eclipse Ti fluorescence microscope) microscope using differential interference contrast (DIC) microscopy at 1000x magnification. Images of recordings were extracted in VLC v3.20 with the snapshoot feature.

### Scanning electron microscopy (SEM)

Approximately 25 mL of the *Hokulea* culture (after ~1 week of growth) was transferred to a large polystyrene petri dish and vapor fixed with OsO_4_ for 10 min at room temperature. Cells were then post-fixed in 1% OsO_4_ for 10 min at room temperature. Fixed cells were transferred onto either a 0.2 μm or 0.4 μm Millipore membrane filters and washed in distilled water once. Fixed cells were then dehydrated using a graded ethanol series (30%, 50%, 70%, 85%, 90%, 95%) and further washed in 100% ethanol three times. Membranes containing fixed and dehydrated cells were dried with CO_2_ using a critical point dryer (UBC Bioimaging Facility). Stubs containing dried membranes were mounted on aluminum SEM stubs and sputter coated with 2 nm Au/Pd. Stubs containing cells were viewed using a Zeiss XB350 Crossbeam scanning electron microscope (SEM) at the UBC Bioimaging Facility. This protocol is adapted from Leander and Farmer (2000).

### Single-cell amplification and sequencing

Individual cells were manually isolated and washed 2**-**4 times using fresh DY-V media. Cells were placed in 0.2 mL PCR strip tubes. The REPLI-g Advance Single Cell Kit (Qiagen) was used to obtain single-cell amplified genome (SAG) following the manufacturer’s protocol. The resulting SAGs were then quantified using a Qubit BR DNA assay (Invitrogen). These SAGs were used as input for the Illumina Nextera DNA Flex Library Prep kit (using half reactions) and then sequenced on an Illumina NovaSeq X series instrument with paired end 2×150 bp reads (performed by Psomagen psomagen.com).

### Single-cell genome assembly

Raw reads were quality trimmed using fastp v.0.23.2 (S. Chen et al. 2018). Trimmed paired reads were use as an input for SPAdes version 4.0.0 (Prjibelski et al. 2020) and assembled under single-cell mode with default parameters, which takes multiple displacement amplification bias into account (Záhonová et al. 2021; Prjibelski et al. 2020; Spits et al. 2006). Contamination was assessed using Blobtoolkit v4.3.5 (Challis et al. 2020) and filtered out via seqkit (Shen et al., 2016). The completeness score for each assembly was assessed with the eukaryota_odb10 database in BUSCO 5.8.0 (Manni et al. 2021). A co-assembly of all SAGs was then performed following the pipeline above and used for all following downstream analyses.

### Phylogenetic reconstruction

SSU rDNA sequences were extracted from the SAG co-assembly using barrnap version 0.9 (https://github.com/tseemann/barrnap/), aligned with a previously published data set (Lax et al., 2020) using MAFFT E-INS-I version 7.481 (Katoh et al., 2013), manually corrected, and then trimmed using trimAl version 1.2rev59 (Capella-Gutiérrez et al., 2009) (-gt 0.87 -st 0.001). A Maximum-likelihood (ML) tree of the final alignment of 239 taxa with 1,811 positions was estimated with RAxML-NG version 1.1.0 (Kozlov et al., 2019) under the GTR + GAMMA model with 1,000 non-parametric bootstrap replicates (Figure 3 and Figure S1).

### Phylogenomic analyses

A multi-gene phylogeny was reconstructed using a previously published 19-gene dataset (Lax et al. 2021). Out of these 19 proteins, we identified 16 in our co-assembly (Table S3). These 16 protein coding genes were extracted manually using a dedicated page on the SAGdb database (https://evocellbio.com/SAGdb/hokulea/), aligned using MUSCLE (Edgar 2004), adjusted using Mesquite v3.81 (Maddison, 2023), and then trimmed using trimAl version 1.2rev59 (Capella-Gutiérrez et al., 2009). Single gene trees were calculated using IQ-Tree2 version 2.3.6 (Minh et al. 2020) and visualized using FigTree v1.4.4 (https://tree.bio.ed.ac.uk/software/figtree/). After our single gene tree screening, and removal of paralogs, contaminant and otherwise aberrant sequences, our final concatenated alignment contained 64 taxa with 7,374 positions. We estimated a multi-gene tree with this concatenation under the LG+C10+F+G4 site-heterogeneous mixture model and 1,000 Ultrafast Bootstraps (UFB) using IQ-Tree2 version 2.3.6 (Mihn et al., 2013). The resulting tree was used as a guide tree to run a posterior mean site frequency analysis (PMSF) under the same model with 200 non-parametric bootstraps (Figure 4 and Figure S2).

## RESULTS & DISCUSSION

### *Hokulea waialensis* n. gen. et sp. is a novel freshwater genus from Hawaii with a streamlined ‘ploeotid’ morphology

We used DIC light microscopy to investigate observable morphological features of a novel ‘ploeotid’ isolated from rock scrapings from Hawaii. Our observations showed that this novel species has all the clade-defining traits of euglenids in general, and several characteristics of ‘ploeotids’, such as 12 pellicle strips, a rigid cell body, a feeding apparatus visible by light microscopy, an anterior flagellum used for feeding, and a posterior flagellum used for gliding (Lax et al., 2019). Cells can be seen ‘tipping’ their body upwards in what could be a characteristic feeding behavior also seen in several other ploeotid taxa like *Ploeotia vitrea* and *Serpenomonas costata* (Movie S3; personal comm. G. Lax). *H. waialensis* cells are colorless, rigid, and oval-shaped with a length of 11.3 μm ± 0.97 μm and width of 5.33 μm ± 0.69 μm (Figure 1A-C). Cells have two flagella originating from the flagellar pocket (Figure 1A-C, Figure 2C). Both flagella differ in thickness and length, with the posterior flagellum being slightly thicker than the anterior flagellum. The anterior flagellum is 5.31 µm ± 1.01 µm long (0.4x cell length) and exhibits a near-constant whipping motion (Movie S1-2). The cell glides using the posterior flagellum, which is 15.50 µm ± 3.41 µm long (1.3x cell length, Figure 1A).

**Figure 1.**
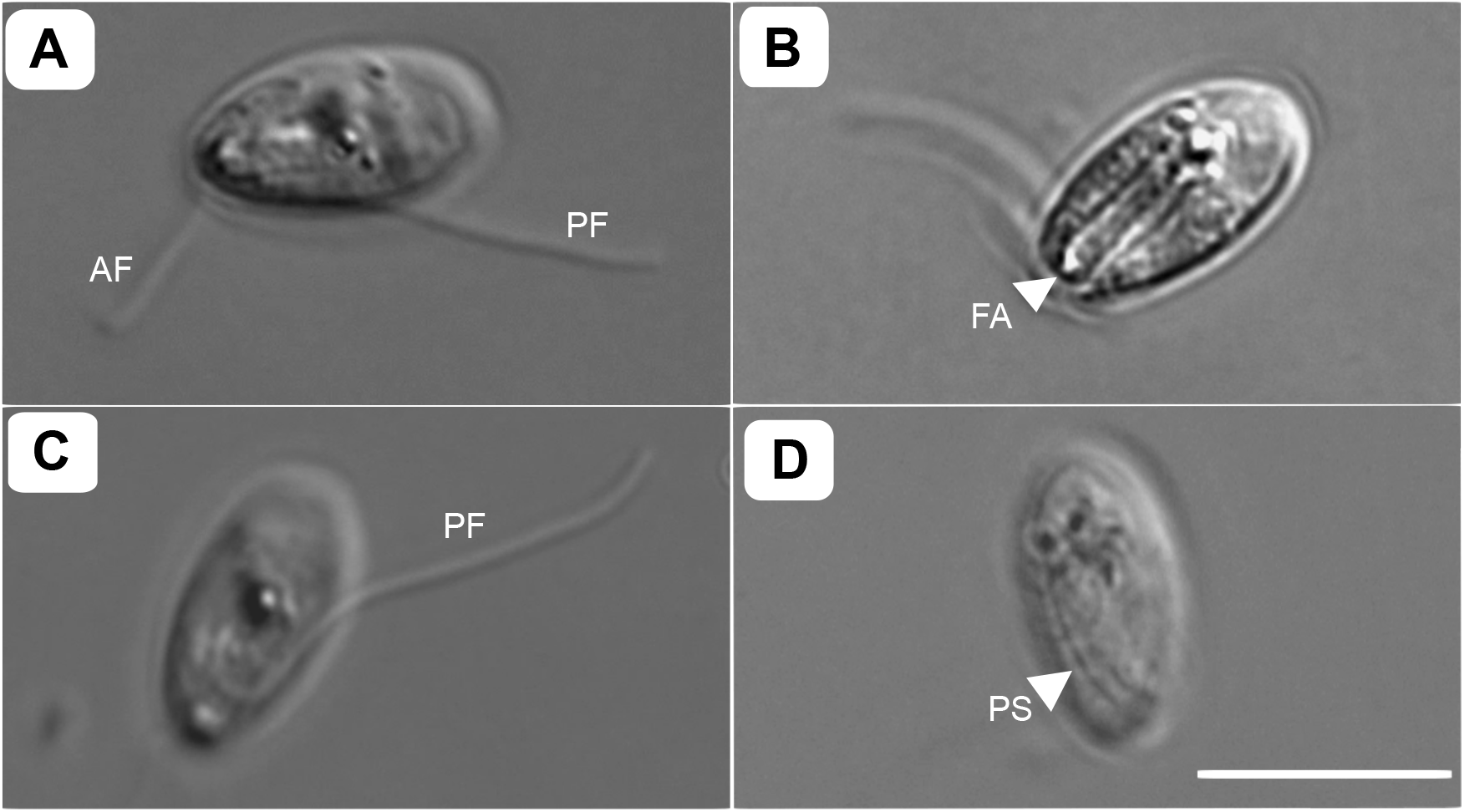
Differential Interference Contrast (DIC) microscopy of *Hokulea waialensis*. (A) DIC image showing the anterior flagellum (AF) and posterior flagellum (PF). (B-C) feeding apparatus (FA), consisting of two robust rods oriented in parallel with the cell (D) pellicle strips (PS). All images are at the same scale as (D), with the scale bar being 10 µm.

**Figure 2.**
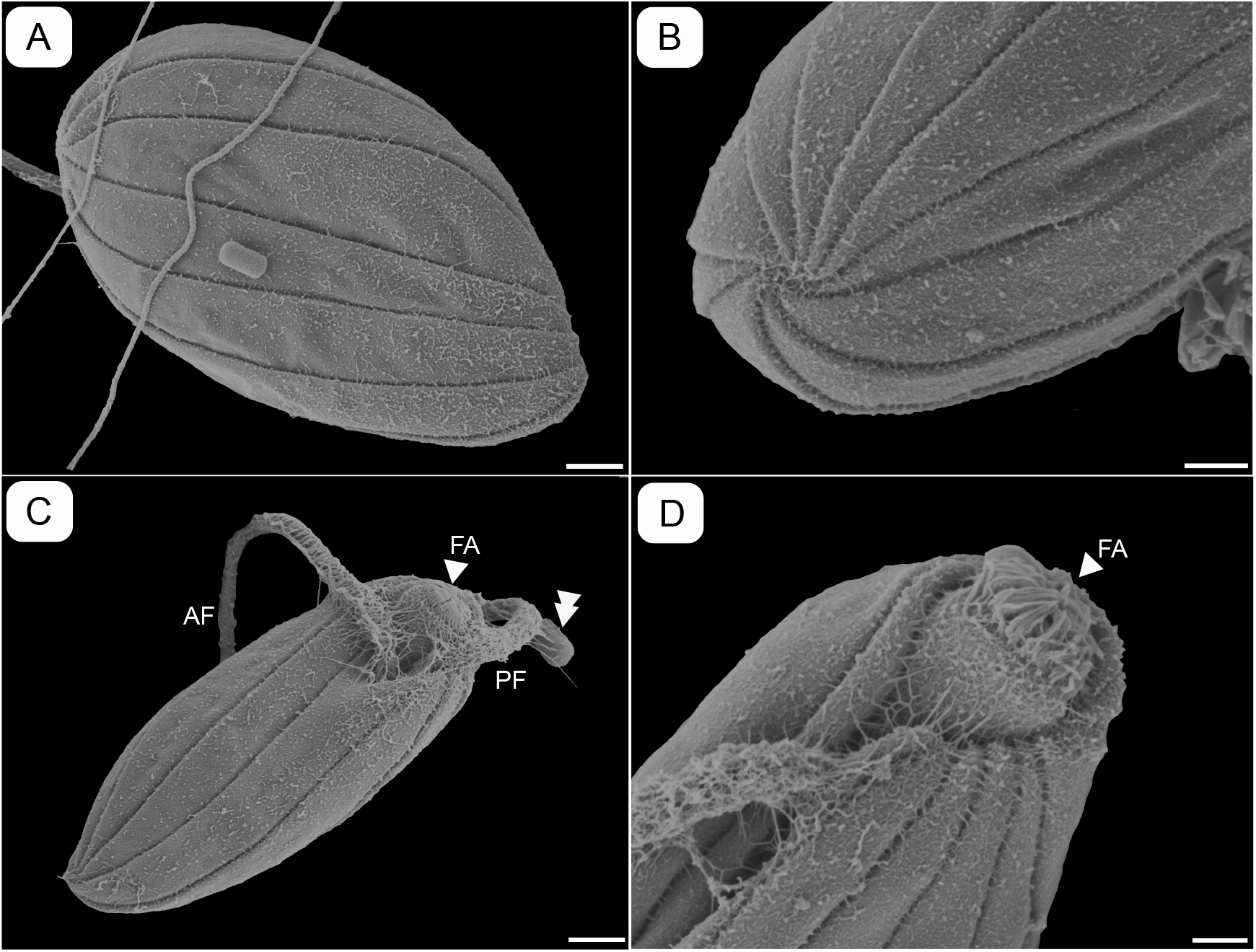
Scanning Electron Microscopy (SEM) of *Hokulea waialensis*. (A) Dorsal view of a cell showing helically arranged pellicle strips. (B) A view of a cell showing ten pellicle strips meeting at the posterior end. (C) Ventral view of a cell showing the anterior flagellum (AF) and the posterior flagellum (PF) emerging from the flagellar pocket and a bacterial cell attached to the PF marked by two arrows. (D) Higher magnification view of the ventral side of a cell showing the feeding apparatus (FA) protruding from the anterior end. Scale bars are 1 µm for A and C, 200 nm for B, and 500 nm for D.

**Figure 3.**
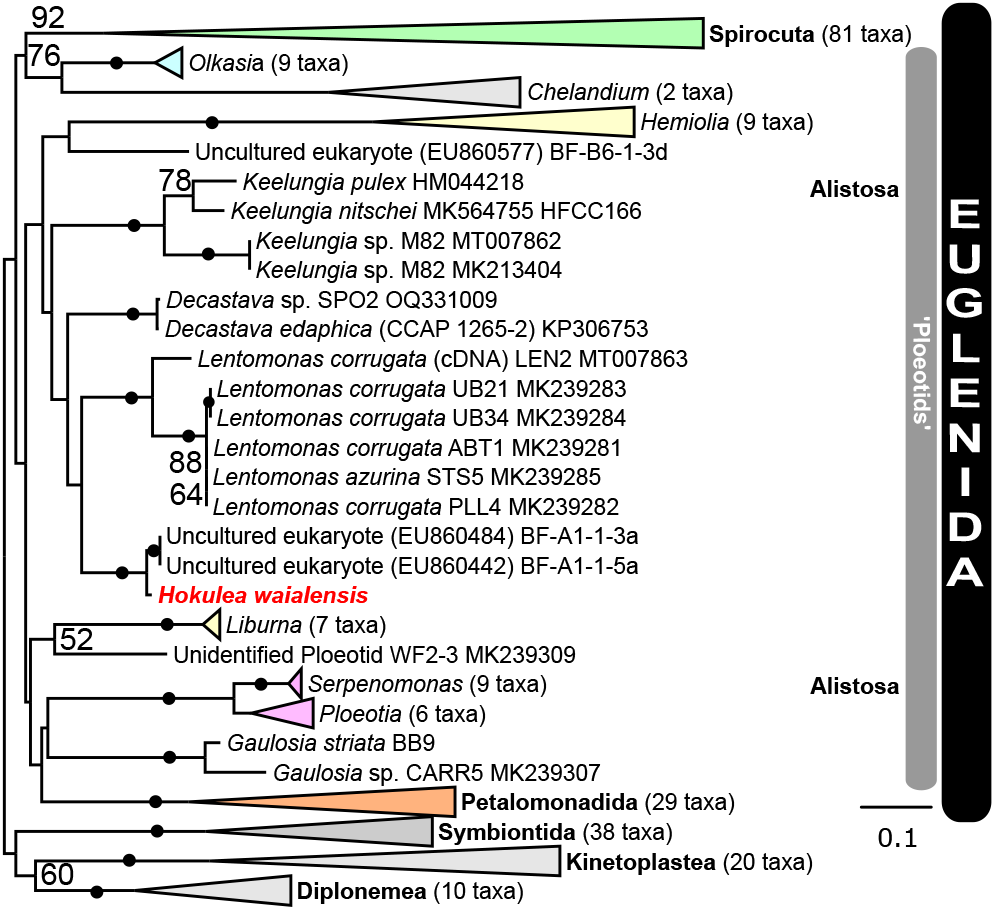
Small Subunit (SSU) rDNA phylogeny showing the position of *Hokulea waialensis* within the Euglenida. The Maximum-likelihood tree was estimated under the GTR + Gamma model with 1,000 nonparametric bootstrap replicates, with a focus on alistosan euglenids, and rooted on outgroups Symbiontida, Diplonemea and Kinetoplastea. *Hokulea waialensis* is marked in red and bold font. Bootstrap values under 50% are not shown, and nodes with 100% support are marked by a black circle. For a version displaying all clades in full, see Supplemental Figure S1.

Previously described ‘ploeotids’ have 10-12 pellicle strips (Leander 2004; Leander et al. 2007, 2017; Lax et al 2019, Triemer 1896). Using scanning electron microscopy (SEM), we clearly show that *H. waialensis* has 10 pellicle strips arranged in a slight helical twist along the longitudinal axis of the cell, all meeting at the posterior tip of the cell (Figure 2 A-B). This differs from some other ploeotids, where the pellicle comprises strictly longitudinally arranged pellicle strips (e.g., *Keelungia pulex, K. nitschei*) (Chan et al., 2014; Schoenle et al. 2019), but is similar to *Ploeotia costaversata* (Schoenle et al. 2019). Unlike other ploeotids whose pellicle strips are either heterogeneous (e.g., narrow, recessed strips alternating with wider strips in *Serpenomonas costata*; (Triemer 1986; Lax et al. 2019) or just projecting outwards like ‘keels’ (e.g., *Ploeotia vitrea, P. oblonga, Olkasia polycarbonata*; Lax et al. 2019), *H. waialensis* contains shallow pellicle grooves (Figure 2 A-C) similar to those of *P. costaversata* and *K. pulex* (Schoenle et al. 2019).

A hook-shaped feeding apparatus can be seen in *H. waialensis* (Figure 1B), which is another prominent feature among ‘ploeotid’ species (Lax et al., 2023). The *H. waialensis* feeding apparatus appears to be morphologically different from other ploeotids having hair-like filaments (Figure 2 C-D), where the lip structure or cap (in the case of *Entosiphon sulcatum* and *Olkasia polycarbonata)* of other ‘ploeotids’ seems to be (Chan et al. 2013; Cavalier-Smith et al. 2016; Lax et al., 2019; Schoenle et al. 2019; Triemer and Fritz 1987; Triemer 1986). It is unclear whether these filaments represent the cap or are situated on top of an existing cap (see Figure 2D). Nonetheless, this is a unique ultrastructural trait that has not been documented in the ultrastructural descriptions of other ploeotid species thus far. The organization and placement of the feeding apparatus with respect to the vestibular opening (i.e., the invagination where both the flagellar and feeding pockets are connected) appears to vary among cells within the same prepared SEM sample. Some cells, particularly in those that lack flagella, the vestibular opening appears reduced and the feeding apparatus appears to be continuous with a ventral pellicle strip (Figure S3 A, B), consistent with other ploeotids like *Lentomonas corrugata* (Triemer 1986; Farmer and Triemer 1994). Contrastingly, in some cells, the vestibular opening appears wider, and the feeding apparatus extends deeper towards the dorsal side of the cell (Figure 2 C, D; Figure S3 C, D). It is unclear if this difference in morphology represents variation among individual cells associated with feeding, or if this is representative of the condition of the cells (e.g., cells that lost their flagella during SEM preparation and are damaged). A detailed ultrastructural investigation of *H. waialensis* using transmission electron microscopy (TEM) would be an exciting avenue of further research to improve our understanding of how the feeding apparatus is organized within the cell. Other notable details observed in *H. waialensis* include a distinct ‘fuzz’ or putative glycocalyx (i.e., extracellular layer) covering the surface of the cell. (Figure 2, Figure S3) (Leander et al. 2017; Lee and Simpson 2014). It is possible that this ‘fuzz’ could also represent a fixation artifact or is specific to the conditions of this culture.

SEM also demonstrates fine hairs covering both flagella that seem to be also thickest around the flagellar pocket (Figure 2 C-D; Figure S3 D). This material can be seen sticking to bacterial cells and may facilitate feeding (Figure 2 C; Figure S3 D). These structures likely represent a combination of mastigonemes on the flagella and tomentum associated with the vestibular opening, similar to that described in other ploeotids like *O. polycarbonata, P. oblonga*, and *S. costata* (Palka et al. 2025; Lax et al. 2021). Given that some heterotrophic euglenids (i.e., *Teloprocta scaphurum*) feed by pulling their prey into the feeding apparatus with their anterior flagellum (Breglia et al. 2013), both the hair-like filaments on the feeding apparatus seen on the SEMs (Figure 2 C-D) and the thread-like material on the flagella could perhaps facilitate feeding in ‘ploeotids’ (Movie S3). Further ultrastructural analysis using TEM is required to compare these hair-like filaments to the ultrastructure traits of other euglenids associated with feeding.

### Single amplified genome of *H. waialensis* provides a baseline for future comparative genomic investigations

We isolated, genome-amplified, and sequenced five cells from a mono-eukaryotic culture via single cell isolation (see Methods). We also co-assembled these SAGs to improve our assembly coverage. To assess assembly quality, we produced blobplots of each individual assembly via Blobtoolkit v4.3.5 (Challis et al. 2020) (Figure S3) from which bacterial contigs and other eukaryotic contaminants were eliminated. We then generated a final co-assembly by re-assembling the filtered contigs. The individual SAGs and the co-assembly are accessible at https://doi.org/10.6084/m9.figshare.29386478.v1. We assessed genome completeness using BUSCO (Manni et al., 2021) in which our individual assemblies ranged from 11% to 33% completeness (Table S1). Our co-assembly had a BUSCO score of 58% (complete + fragmented, Table S1). This is reasonably complete when compared to the *Euglena gracilis* genome at 73% (Chen et al. 2024), which is considered the most complete euglenid genome to date. Though several investigations have generated single amplified transcriptomes (SATs), this is the first that generated SAGs from a phagotrophic euglenid via single cell isolation. Previous SAG studies have either involved flow sorting (Záhonová et al. 2021; Wideman et al. 2019), single cell isolation of other groups of euglenozoans (e.g., eupelagonemids (Gawryluk et al. 2016), or genome-amplifying phagotrophic euglenids for the purpose of PCR-amplifying the SSU rDNA (Lax et al. 2019; Lax and Simpson 2020).

### *Hokulea* is a novel genus branching within Alistosa, closely related to *Decastava* and Lentomonas

The SSU rDNA sequences from *H. waialensis* were extracted from the SAGs of the co-assembly via barrnap, and a phylogenetic tree was inferred using a previously published dataset (Lax et al. 2019). Like analyses in other studies, our SSU rDNA phylogeny was poorly resolved in its deeper nodes, though *H. waialensis* branched close to both *Decastava* and *Lentomonas*, forming a clade with two environmental sequences, BF-A1-1-3a and BF-A1-1-5a (Figure 4; Figure S2) with maximum bootstrap support (100%). We extracted 16 protein coding genes to reconstruct a multigene phylogeny based on a previously published alignment (Lax et al. 2023). Our phylogenetic analyses are in line with this most recent multigene phylogenies of euglenids (Lax et al. 2023): Euglenids form a monophyletic group with high support in all our analysis (100/96% BS/PMSF, Figure 4, Figure S2), and the major groups of euglenids were recovered with full support. These included the species-rich petalomonads as the deepest branch (100% BS/PMSF, Figure 4, Figure S2) and Spirocuta, which includes well-studied phototrophic euglenids (100% BS/PMSF).

**Figure 4.**
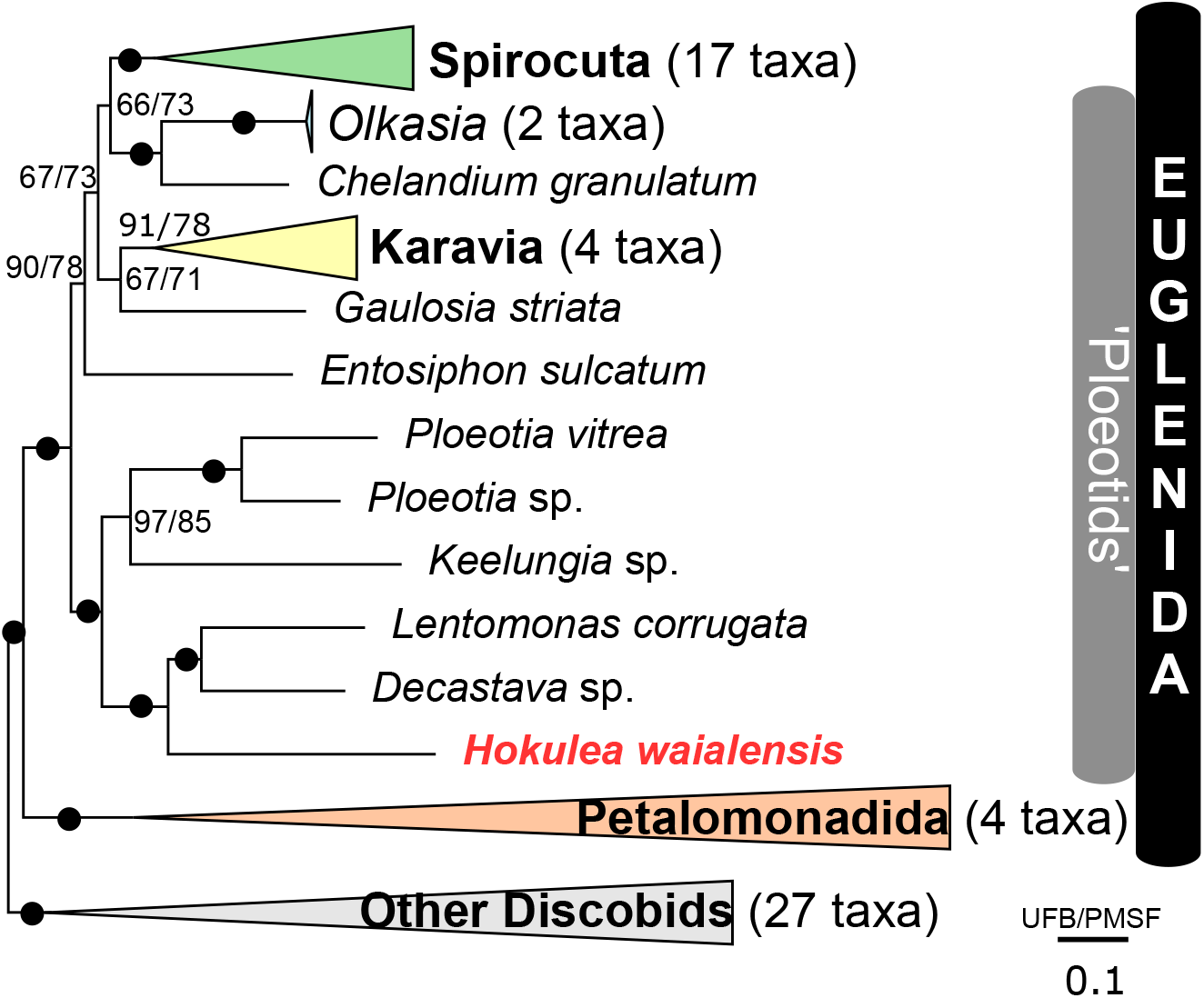
Multigene phylogeny of 64 discobid taxa using 16 genes, with a focus on euglenids (7374 total amino acid sites). Estimated under Maximum-likelihood and the LG+C10+F+G4 model with 1000 Ultrafast bootstraps (UFB) and 200 non-parametric bootstraps (PMSF). Rooted on other discobids. *Hokulea waialensis* is marked in red and bold font. Nodes that received >95% support from both ultrafast bootstrap (UFB) and posterior mean site frequency (PMSF) analyses are marked with a black circle. For a version displaying all clades in full, see Supplemental Figure S2).

The group Alistosa was also recovered with full support (100% BS/PMSF, Figure 4; Figure S2), with *H. waialensis* branching as sister to a clade comprising *Decastava* and *Lentomonas* (100% BS/PMSF). Although there is still missing multigene data from other ‘ploeotids’ in key phylogenomic positions, such as *Serpenomonas* and uncharacterized taxa such as WF2-3 (Lax et al. 2021), our data confirm previous taxonomic assignments. Our high completion estimates of the *H. waialensis* SAGs suggest that genomes might be more easily generated for these taxa and should be considered in future investigations.

## Conclusions

The phylogenetic placement of *H. waialensis* within the Alistosa is consistent with the morphological features observed here, including the presence of 10 pellicles strips and the hook-shaped feeding apparatus (Figure 1C). A potential distinguishing feature for this species is the presence of fibrous material in and around the feeding apparatus visible with SEM (Figure 2 C-D; Figure S3 A-C), which has not been previously reported in other ploeotids. Although ploeotids comprise much of the phylogenetic diversity of phagotrophic euglenids, most described species are seemingly from marine environments (Lee and Patterson 2000; Larsen and Patterson 1990; Lax et al. 2019; Lax et al. 2023) except for *Entosiphon* (freshwater) and *Decastava* (soil) (Lax et al. 2023). The lack of freshwater ploeotids species could be due to a sampling bias on marine environments or simply because ploeotids are not particularly common in freshwater environments. Furthermore, even if environmental sequences can provide evidence of hidden diversity in these poorly studied environments, the primer bias for euglenids make these studies inadequate for understanding euglenid diversity (Lax et. al., 2019, Lax et. al., 2020, Lax et. al., 2023), because the vast majority of euglenid diversity is missed. *H. waialensis* represents part of this missing freshwater ploeotid diversity branching with two other freshwater sequences in our SSU rDNA phylogeny, further promoting its uniqueness and showcasing the importance for further taxonomic sampling and characterization of ploeotids from freshwater environments. Sampling in non-marine environments will ultimately help us better understand the evolutionary history of phagotrophic euglenids specifically, and euglenids in general. Additional SAGs from other ploeotid taxa would also help resolve branching patterns and allow us to make inferences about character evolution in euglenozoans.

## Taxonomic descriptions

### Hokulea

gen. nov. Rodriguez Ruiz, Palka, Lax, Jirsová, Fuggiti, Poh, Leander, and Wideman (ICZN).

### Taxonomic summary

Eukaryota; Discoba; Euglenozoa; Euglenida.

### Diagnosis

Free living, rigid, biflagellated heterotrophic euglenid with an oblong cell shape. Cell body slightly flattened ventrally. Anterior flagellum ~0.4x cell length and posterior flagellum 1.3x cell length. Slow gliding speed with hook-shaped feeding apparatus, and 10 pellicle strips, only some of which can be readily seen with light microscopy.

### Type species

*Hokulea waialensis* Rodriguez Ruiz, Palka, Lax, Jirsová, Fuggiti, Poh, Leander, and Wideman

### Etymology

From ‘Hōkulea’ (Hawaiian, singular) a traditional Polynesian boat from Hawaii, also known as the star of Arcturus. It is a symbol of hope for the Hawaiian indigenous people.

### ZooBank Accession

LSID urn:lsid:zoobank.org:act:7AA6D9B0-1C9B-4540-BD36-3CD0E3422B7E

### Hokulea waialensis

sp. nov. Rodriguez Ruiz, Palka, Lax, Jirsová, Fuggiti, Poh, Leander, and Wideman

(ICZN).

### Diagnosis

Oblong cell shape with slightly rounded posterior, 11.3 μm long and 4.9 μm wide ventrally flattened.

### Type material

Cell imaged in Figure 1 and Figure 2

### Type habitat

Freshwater oxic sediment

### Type locality

Wai’ale Falls, Wailuku River, Hilo, Hawaii. (19.715° N, −155.138° W)

### Etymology

From ‘Waialensis’ (Hawaiian) owing to the place to which the sample was collected (Wai’ale Falls)

### Gene sequence

The SSU rDNA sequence extracted from the single-cell genome assembly of *H. waialensis* is deposited under GenBank Accession number PQ611217.

### ZooBank Accession

LSID urn:lsid:zoobank.org:act:7AA6D9B0-1C9B-4540-BD36-3CD0E3422B7E

## Acknowledgements

This work was funded by National Science Foundation grants to JGW (2119963 and 2405455) and a grant to BSL from the National Sciences and Engineering Research Council of Canada (NSERC 2025-04268).

## References

Andersen, Robert A., ed. 2009. Algal Culturing Techniques. Nachdr. Amsterdam Heidelberg: Elsevier Academic Press.

Breglia, Susana A., Claudio H. Slamovits, and Brian S. Leander. 2007. “Phylogeny of Phagotrophic Euglenids (Euglenozoa) as Inferred from Hsp90 Gene Sequences.” Journal of Eukaryotic Microbiology 54 (1): 86–92. 10.1111/j.1550-7408.2006.00233.x.

Breglia, Susana A., Naoji Yubuki, and Brian S. Leander. 2013. “Ultrastructure and Molecular Phylogenetic Position of Heteronema Scaphurum : A Eukaryovorous Euglenid with a Cytoproct.” Journal of Eukaryotic Microbiology 60 (2): 107–20. 10.1111/jeu.12014.

Camacho, Christiam, George Coulouris, Vahram Avagyan, Ning Ma, Jason Papadopoulos, Kevin Bealer, and Thomas L Madden. 2009. “BLAST+: Architecture and Applications.” BMC Bioinformatics 10 (1): 421. 10.1186/1471-2105-10-421.

Capella-Gutiérrez, Salvador, José M. Silla-Martínez, and Toni Gabaldón. 2009. “trimAl: A Tool for Automated Alignment Trimming in Large-Scale Phylogenetic Analyses.” Bioinformatics 25 (15): 1972–73. 10.1093/bioinformatics/btp348.

Cavalier-Smith, Thomas. 2017. “Euglenoid Pellicle Morphogenesis and Evolution in Light of Comparative Ultrastructure and Trypanosomatid Biology: Semi-Conservative Microtubule/Strip Duplication, Strip Shaping and Transformation.” European Journal of Protistology 61 (October):137–79. 10.1016/j.ejop.2017.09.002.

Cavalier-Smith, Thomas, Ema E. Chao, and Keith Vickerman. 2016. “New Phagotrophic Euglenoid Species (New Genus Decastava; Scytomonas Saepesedens; Entosiphon Oblongum), Hsp90 Introns, and Putative Euglenoid Hsp90 Pre-mRNA Insertional Editing.” European Journal of Protistology 56 (October):147–70. 10.1016/j.ejop.2016.08.002.

Challis, Richard, Edward Richards, Jeena Rajan, Guy Cochrane, and Mark Blaxter. 2020. “BlobToolKit – Interactive Quality Assessment of Genome Assemblies.”

Chan, Ya-Fan, øjvind Moestrup, and Jeng Chang. 2013. “On Keelungia Pulex Nov. Gen. et Nov. Sp., a Heterotrophic Euglenoid Flagellate That Lacks Pellicular Plates (Euglenophyceae, Euglenida).” European Journal of Protistology 49 (1): 15–31. 10.1016/j.ejop.2012.04.003.

Chen, Shifu, Yanqing Zhou, Yaru Chen, and Jia Gu. 2018. “Fastp: An Ultra-Fast All-in-One FASTQ Preprocessor.” Bioinformatics 34 (17): i884–90. 10.1093/bioinformatics/bty560.

Chen, Zixi, Yang Dong, Shengchang Duan, Jiayi He, Huan Qin, Chao Bian, Zhenfan Chen, et al. 2024. “A Chromosome-Level Genome Assembly for the Paramylon-Producing Microalga Euglena gracilis.” Scientific Data 11 (1): 780. 10.1038/s41597-024-03404-y.

Edgar, Robert C. 2004. “MUSCLE: A Multiple Sequence Alignment Method with Reduced Time and Space Complexity.” BMC Bioinformatics 5 (1): 113. 10.1186/1471-2105-5-113.

Esson, Heather J. and Leander Brian S. 2008. Novel pellicle surface patterns on Euglena obtusa Schmitz (Euglenophyta), a euglenophyte from a benthic marine environment: Implications for pellicle development and evolution. J. Phycol. 44: 132–141. https://onlinelibrary.wiley.com/doi/abs/10.1111/j.1529-8817.2007.00447.x

Farmer, Mark A., and Richard E. Triemer. 1994. “An Ultrastructural Study of Lentomonas Applanatum (Preisig) N. G. (Euglenida).” Journal of Eukaryotic Microbiology 41 (2): 112–19. 10.1111/j.1550-7408.1994.tb01482.x.

Gawryluk, Ryan M.R., Javier Del Campo, Noriko Okamoto, Jürgen F.H. Strassert, Julius Lukeš, Thomas A. Richards, Alexandra Z. Worden, Alyson E. Santoro, and Patrick J. Keeling. 2016. “Morphological Identification and Single-Cell Genomics of Marine Diplonemids.” Current Biology 26 (22): 3053–59. 10.1016/j.cub.2016.09.013.

Kostygov, Alexei Y., Anna Karnkowska, Jan Votýpka, Daria Tashyreva, Kacper Maciszewski, Vyacheslav Yurchenko, and Julius Lukeš. 2021. “Euglenozoa: Taxonomy, Diversity and Ecology, Symbioses and Viruses.” Open Biology 11 (3): 200407. 10.1098/rsob.200407.

Kozlov, Alexey M, Diego Darriba, Tomáš Flouri, Benoit Morel, and Alexandros Stamatakis. 2019. “RAxML-NG: A Fast, Scalable and User-Friendly Tool for Maximum Likelihood Phylogenetic Inference.” Edited by Jonathan Wren. Bioinformatics 35 (21): 4453–55. 10.1093/bioinformatics/btz305.

Larsen, Jacob, and David J. Patterson. 1990. “Some Flagellates (Protista) from Tropical Marine Sediments.” Journal of Natural History 24 (4): 801–937. 10.1080/00222939000770571.

Lax, G., M. Kolisko, Y. Eglit, W.J. Lee, N. Yubuki, A. Karnkowska, B.S. Leander, G. Burger, P.J. Keeling, and A.G.B. Simpson. 2021. “Multigene Phylogenetics of Euglenids Based on Single-Cell Transcriptomics of Diverse Phagotrophs.” Molecular Phylogenetics and Evolution 159 (June):107088. 10.1016/j.ympev.2021.107088.

Lax, Gordon, Anna Cho, and Patrick J. Keeling. 2023. “Phylogenomics of Novel Ploeotid Taxa Contribute to the Backbone of the Euglenid Tree.” Journal of Eukaryotic Microbiology 70 (4): e12973. 10.1111/jeu.12973.

Lax, Gordon, and Patrick J. Keeling. 2023. “Molecular Phylogenetics of Sessile Dolium Sedentarium, a Petalomonad Euglenid.” Journal of Eukaryotic Microbiology 70 (5): e12991. 10.1111/jeu.12991.

Lax, Gordon, Won Je Lee, Yana Eglit, and Alastair Simpson. 2019. “Ploeotids Represent Much of the Phylogenetic Diversity of Euglenids.” Protist 170 (2): 233–57. 10.1016/j.protis.2019.03.001.

Lax, Gordon, and Alastair G.B. Simpson. 2020. “The Molecular Diversity of Phagotrophic Euglenids Examined Using Single-Cell Methods.” Protist 171 (5): 125757. 10.1016/j.protis.2020.125757.

Leander, Brian S. 2004. Did trypanosomatid parasites have photosynthetic ancestors? Trends Microbiol. 12:251–258. https://www.sciencedirect.com/science/article/abs/pii/S0966842X04000927

Leander, Brian S., and Mark A. Farmer. 2000. “Comparative Morphology of the Euglenid Pellicle. I. Patterns of Strips and Pores.” Journal of Eukaryotic Microbiology 47 (5): 469–79. 10.1111/j.1550-7408.2000.tb00076.x.

Leander, Brian S., and Mark A. Farmer. 2001. “Comparative Morphology of the Euglenid Pellicle. II. Diversity of Strip Substructure.” Journal of Eukaryotic Microbiology 48 (2): 202–17. 10.1111/j.1550-7408.2001.tb00304.x.

Leander, Brian S., Esson Heather J. and Breglia Susana A. 2007. Macroevolution of complex cytoskeletal systems in euglenids. BioEssays. 29: 987–1000. https://onlinelibrary.wiley.com/doi/abs/10.1002/bies.20645

Leander, Brian S., Gordon Lax, Anna Karnkowska, and Alastair G. B. Simpson. 2017. “Euglenida.” In Handbook of the Protists, edited by John M. Archibald, Alastair G.B. Simpson, Claudio H. Slamovits, Lynn Margulis, Michael Melkonian, David J. Chapman, and John O. Corliss, 1–42. Cham: Springer International Publishing. 10.1007/978-3-319-32669-6_13-1.

Leander, Brian S., Richard E. Triemer, and Mark A. Farmer. 2001. “Character Evolution in Heterotrophic Euglenids.” European Journal of Protistology 37 (3): 337–56. 10.1078/0932-4739-00842.

Li, Heng. 2018. “Minimap2: Pairwise Alignment for Nucleotide Sequences.” Edited by Inanc Birol. Bioinformatics 34 (18): 3094–3100. 10.1093/bioinformatics/bty191.

Maddison, W.P., and D.R. Maddison. 2023. “Mesquite: A Modular System for Evolutionary Analysis.”

Manni, Mosè, Matthew R. Berkeley, Mathieu Seppey, and Evgeny M. Zdobnov. 2021. “BUSCO: Assessing Genomic Data Quality and Beyond.” Current Protocols 1 (12): e323. 10.1002/cpz1.323.

Minh, B. Q., M. A. T. Nguyen, and A. Von Haeseler. 2013. “Ultrafast Approximation for Phylogenetic Bootstrap.” Molecular Biology and Evolution 30 (5): 1188–95. 10.1093/molbev/mst024.

Minh, Bui Quang, Heiko A Schmidt, Olga Chernomor, Dominik Schrempf, Michael D Woodhams, Arndt Von Haeseler, and Robert Lanfear. 2020. “IQ-TREE 2: New Models and Efficient Methods for Phylogenetic Inference in the Genomic Era.” Edited by Emma Teeling. Molecular Biology and Evolution 37 (5): 1530–34. 10.1093/molbev/msaa015.

Palka, Maia V., Regine Claire Manglicmot, Gordon Lax, Kevin C. Wakeman, and Brian S. Leander. 2025. “Ultrastructure of Olkasia Polycarbonata (Euglenozoa, Euglenida) Demonstrates Cytoskeletal Innovations Associated with the Feeding and Flagellar Apparatuses.” Journal of Eukaryotic Microbiology 72 (1). 10.1111/jeu.13074.

Prjibelski, Andrey, Dmitry Antipov, Dmitry Meleshko, Alla Lapidus, and Anton Korobeynikov. 2020. “Using SPAdes De Novo Assembler.” Current Protocols in Bioinformatics 70 (1): e102. 10.1002/cpbi.102.

Schoenle, Alexandra, Suzana Živaljić, Dennis Prausse, Janine Voß, Kirsten Jakobsen, and Hartmut Arndt. 2019. “New Phagotrophic Euglenids from Deep Sea and Surface Waters of the Atlantic Ocean (Keelungia Nitschei, Petalomonas Acorensis, Ploeotia Costaversata).” European Journal of Protistology 69 (June):102–16. 10.1016/j.ejop.2019.02.007.

Shen, Wei, Shuai Le, Yan Li, and Fuquan Hu. 2016. “SeqKit: A Cross-Platform and Ultrafast Toolkit for FASTA/Q File Manipulation.” Edited by Quan Zou. PLOS ONE 11 (10): e0163962. 10.1371/journal.pone.0163962.

Strother, Paul K., Wilson A. Taylor, Bas Van De Schootbrugge, Brian S. Leander, and Charles H. Wellman. 2020. “Pellicle Ultrastructure Demonstrates That Moyeria Is a Fossil Euglenid.” Palynology 44 (3): 461–71. 10.1080/01916122.2019.1625457.

Triemer, Richard E. 1986. “Light and Electron Microscopic Description of a Colorless Euglenoid, Serpenomonas Costata n. g., n. Sp. 1.” The Journal of Protozoology 33 (3): 412–15. 10.1111/j.1550-7408.1986.tb05632.x.

Triemer, Richard E., and Lawrence Fritz. 1987. “Structure and Operation of the Feeding Apparatus in a Colorless Euglenoid, Entosiphon Sulcatum 1.” The Journal of Protozoology 34 (1): 39–47. 10.1111/j.1550-7408.1987.tb03129.x.

Valster, Rinske M., Bart A. Wullings, Geo Bakker, Hauke Smidt, and Dick Van Der Kooij. 2009. “Free-Living Protozoa in Two Unchlorinated Drinking Water Supplies, Identified by Phylogenic Analysis of 18S rRNA Gene Sequences.” Applied and Environmental Microbiology 75 (14): 4736–46. 10.1128/AEM.02629-08.

Wideman, Jeremy G., Gordon Lax, Guy Leonard, David S. Milner, Raquel Rodríguez-Martínez, Alastair G. B. Simpson, and Thomas A. Richards. 2019. “A Single-Cell Genome Reveals Diplonemid-like Ancestry of Kinetoplastid Mitochondrial Gene Structure.” Philosophical Transactions of the Royal Society B: Biological Sciences 374 (1786): 20190100. 10.1098/rstb.2019.0100.

Yubuki, Naoji and Leander Brian S. 2012. Reconciling the bizarre inheritance of microtubules in complex (euglenid) microeukaryotes. Protoplasma. 249: 859–869. https://link.springer.com/article/10.1007/s00709-011-0340-z

Záhonová, Kristína, Gordon Lax, Savar D. Sinha, Guy Leonard, Thomas A. Richards, Julius Lukeš, and Jeremy G. Wideman. 2021. “Single-Cell Genomics Unveils a Canonical Origin of the Diverse Mitochondrial Genomes of Euglenozoans.” BMC Biology 19 (1): 103. 10.1186/s12915-021-01035-y.

